# Sexual transmission of *Anopheles gambiae* densovirus (AgDNV) leads to disseminated infection in mated females

**DOI:** 10.1101/2022.02.10.479983

**Authors:** Kristine L Werling, Rebecca M. Johnson, Hillery C Metz, Jason L Rasgon

## Abstract

**Background:** *Anopheles gambiae* densovirus (AgDNV) is an insect-specific, single-stranded DNA virus that infects *An. gambiae*, the major mosquito species responsible for transmitting malaria parasites throughout Sub-Saharan Africa. AgDNV is a benign virus that is very specific to its mosquito host and therefore has potential to serve as a vector control tool via paratransgenesis (genetic modification of mosquito symbionts) to limit transmission of human pathogens. Prior to being engineered into a control tool, the natural transmission dynamics of AgDNV between *An. gambiae* mosquitoes needs to be fully understood. Additionally, improved knowledge of AgDNV infection in male mosquitoes is needed. In this study, we examine the tissue tropism of AgDNV in the male reproductive tract and investigate both venereal and vertical transmission dynamics of the virus.

**Methods:** *An. gambiae* adult males were infected with AgDNV via micro-injection and reproductive tissues collected and assayed for AgDNV using qPCR. Next, uninfected females were introduced to AgDNV-infected or control males and, after several nights of mating, both the spermatheca and female carcass were assessed for venereally transmitted AgDNV. Finally, F1s from this cross were collected and assayed to quantify vertical transmission of the virus.

**Results:** AgDNV infected the reproductive tract of male mosquitoes, including the testes and male accessory glands (MAGs), without affecting mating rates. AgDNV-infected males venereally transmitted virus to females, and these venereally-infected females developed disseminated infection throughout the body. However, AgDNV was not vertically transmitted to F1s resulting from this cross.

**Conclusions:** Infected male releases could be an effective strategy to introduce AgDNV-based paratransgenic tools into naïve populations of *An. gambiae* females.

## Background

*Anopheles gambiae* densovirus (AgDNV) is an insect specific virus (ISV) that efficiently infects *An. gambiae* mosquitoes (1, 2), one of the major vectors of malaria-causing parasites. This non-enveloped, single-stranded DNA virus belongs to the *Parvoviridae* family and *Brevidensovirus* genus and has a compact genome of ~4.1kb (1). First identified in an *An. gambiae* cell line (Sua5B) (1), AgDNV is very specific to *An. gambiae* mosquitoes; it replicates only minimally in closely related *Anopheles* species and is unable to infect vertebrates (2). Due to this host specificity, ISVs like AgDNV offer a promising new avenue for targeted biological control of individual mosquito species and the pathogens they transmit (3–5). In its natural *An. gambiae* host, AgDNV can infect both sexes and all developmental stages, replicating particularly well in adults post-emergence (6, 7). Notably though, AgDNV is a benign virus that does not cause mortality or even a marked transcriptional response in its mosquito host (8). This is in contrast to several related mosquito-specific densoviruses that infect *Aedes* mosquitoes with high lethality (9–11). While cytotoxic densoviruses can be harnessed to make novel biopesticides (5) – as was done with the *Aedes aegypti* densovirus (AaeDNV) (12) – the benign nature of AgDNV is highly advantageous for another vector control approach known as paratransgenesis, or the genetic modification of symbionts with the goal of decreasing pathogen transmission. Specifically, AgDNV could be genetically modified to encode a specific effector that, upon delivery and expression in its *An. gambiae* host, targets either the mosquito or the pathogens they transmit to ultimately decrease incidence of mosquito-borne diseases like malaria. The potential use of AgDNV for paratransgenesis has previously been demonstrated (1, 7, 13), but for an AgDNV-based tool to be effective in field settings, we first need a complete understanding of AgDNV transmission dynamics between *An. gambiae* hosts. This information is critical for determining the best application and deployment strategy for an AgDNV-based vector control tool.

Previously, it was found that AgDNV can be venereally transmitted from infected males to the spermatheca of females post-mating (6), yet it is not known if venereally transmitted AgDNV remains contained to the spermatheca or if it can also lead to disseminated infection in females. This information is important for determining whether infected male releases could be a viable method for introducing AgDNV-based tools into field populations of female *An. gambiae.* As only female mosquitoes blood feed and transmit disease-causing pathogens, they are the main target of vector control programs, and as such, male mosquitoes can be safely released into a population without directly affecting pathogen transmission. Modified male releases is an approach that has been successful for certain *Aedes aegypti* control programs using either the endosymbiont bacterium *Wolbachia* (14) or transgenic male mosquitoes (15, 16). In addition to venereal transmission, AgDNV can also be vertically transmitted from mother to offspring when the mother is infected during larval stages (1), but it is not yet known if mothers infected venereally can subsequently transmit AgDNV to the next generation. This would reveal if AgDNV-based tools could disseminate trans-generationally into a population following infected male releases.

We also do not know the tissue tropism of AgDNV in male mosquitoes, particularly in their reproductive tissues where infection could impact mating success. Notably, *plugin* is one of the few genes that is differentially expressed in AgDNV-infected mosquitoes (8). It encodes a fundamental component of the male mating plug, a coagulated mass of seminal secretions transferred to females during copulation that is essential for proper sperm storage in females (17). Given that AgDNV influences the expression of this key mating factor, assessing whether AgDNV infection influences male mating rates is also an urgent question, as this could impinge on the success of AgDNV-based tools.

Here we address these questions by examining AgDNV tropism and transmission dynamics in lab infected mosquitoes. We find that AgDNV spreads venereally from infected males to naïve females, such that females develop disseminated infection; however it is not subsequently transmitted vertically to F1 offspring. Our findings demonstrate that AgDNV holds promise as a tool for paratransgenesis in *An. gambiae*, whereby infected male releases could be utilized for dissemination into field populations.

## Results

### AgDNV infects male reproductive tissues without affecting mating rates

To determine the tissue tropism and infection dynamics of AgDNV in the reproductive tract of *An. gambiae* males, we infected young adult males with AgDNV via inter-thoracic microinjections and then assessed viral titers at four timepoints, relative to PBS-injected controls. We found that AgDNV infects both major male reproductive tissues – the testes (Figure 1A) and the male accessory glands (MAGs) (Figure 1B) – as well as the remainder of the body (hereafter referred to as carcass) (Figure 1C).

**Figure 1:**
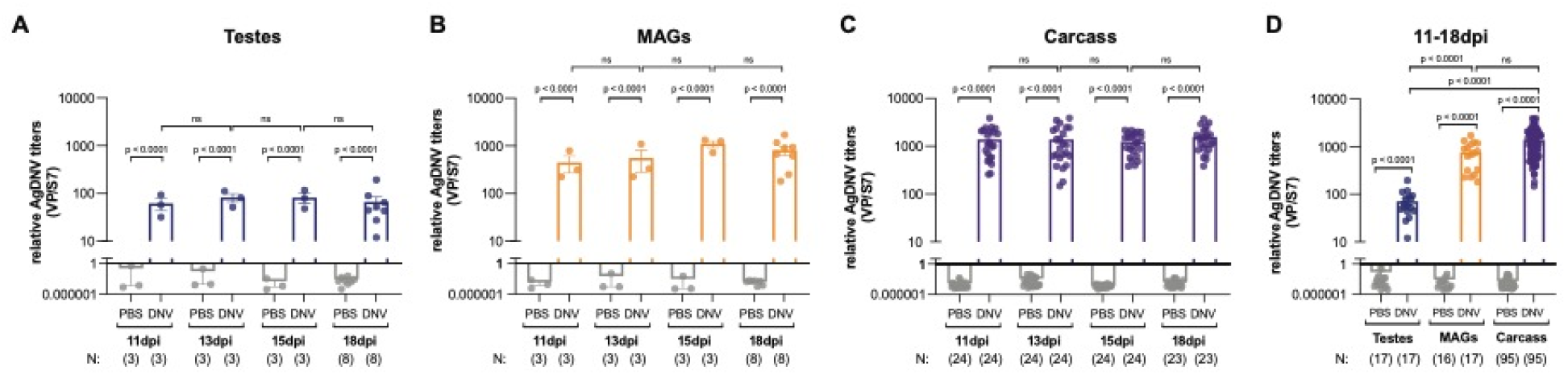
AgDNV infects the reproductive tract of *An. gambiae* males. (**A-C**) Males were injected with either AgDNV (DNV) or PBS (control). At 11-, 13-, 15- and 18-days post-injection (dpi), AgDNV-treated males have infected (**A**) testes (Ln(y) transform, One-way ANOVA, Sidak’s correction), (**B**) MAGs (Ln(y) transform, One-way ANOVA, Sidak’s correction), and (**C**) carcass (Ln(y) transform, Brown-Forsythe/Welch ANOVA, Dunnett’s correction), yet viral titers do not change over time in any tissue. (**D**) Independent of time (including 11-18dpi), AgDNV titers in the MAGs and carcass are significantly higher than those in the testes (Ln(y) transform, Brown-Forsythe/Welch ANOVA, Dunnett’s correction). NS indicates p-value > 0.05. Throughout, for testes and MAGs, each point represents a pool of 12 tissues; for carcass, each point represents an individual carcass with the reproductive tract removed. Viral genomes (VP = viral protein) are normalized to host genomes (S7).

AgDNV was detectable in all tissues at 11-, 13-, 15-, and 18-days post-injection (dpi), yet within each tissue type, relative viral titers (i.e., normalized to host genome) did not differ significantly between time points (Figure 1A-C). Independent of time post-injection, we found significantly higher relative AgDNV titers in the MAGs compared to the testes (Figure 1D), with MAG titers comparable to those of the carcass. In sum, we find that both components of the male reproductive tract are susceptible to infection, but AgDNV likely replicates more efficiently in the MAGs than in the testes.

To assess whether AgDNV infection in male mosquitoes affects their ability to mate, we introduced uninfected, virgin females into cages with either AgDNV- or PBS-injected males. After three nights of mating, females were dissected and mating status was determined by the presence or absence of sperm in the spermatheca. We found no difference in the proportion of mated females between those exposed to AgDNV-injected males and those exposed to PBS-injected males (Table 1), indicating that AgDNV infection does not significantly hinder male mating ability. Additionally, we found no difference in the survival of AgDNV-injected males compared to controls when assessed at 18dpi (Table 2), supporting previous findings that AgDNV infection does not affect longevity of its *An. gambiae* host (8).

**Table 1:** Mating rates.

|  | PBS |  | AgDNV |  |
| --- | --- | --- | --- | --- |
|  | Mated | Unmated | Mated | Unmated |
| P | 29% | 71% | 36% | 64% |
| N | 46 | 111 | 63 | 114 |
| Total | 157 |  | 177 |  |

There is no difference in the mating rate between PBS- and AgDNV-injected males (Fisher's Exact, $p = 0.2432$ ), as determined by the number of inseminated females. $P =$ percent.

**Table 2:** Male survival rates.

|  | PBS |  | AgDNV |  |
| --- | --- | --- | --- | --- |
|  | Alive | Dead | Alive | Dead |
| P | 71% | 29% | 75% | 25% |
| N | 371 | 151 | 428 | 141 |
| Total | 522 |  | 569 |  |

There is no difference in the survival of PBS- and AgDNV-injected males (Fisher's Exact, $p = 0.1322$ ) when assessed at 18 days post-injection. $P =$ percent.

### *An. gambiae* females develop disseminated infection after mating with AgDNV-infected males

Previous studies reported that AgDNV is venereally transmitted from infected males to the spermatheca of mated females (6). To test whether sexually-transmitted AgDNV can also lead to disseminated infection in mated females, we again allowed uninfected, virgin females to mate with either AgDNV- or PBS-injected males. After mating, the females were separated and maintained for 8-14 days before being analyzed. As expected, we detected AgDNV in the spermatheca of females mated to infected males (Figure 2A). We also found significant levels of AgDNV in the female carcass (Figure 2B). Although AgDNV titers were variable, an estimated 90% of mated females (n = 45) developed disseminated infection from venereally transmitted AgDNV (see Methods).

**Figure 2:**
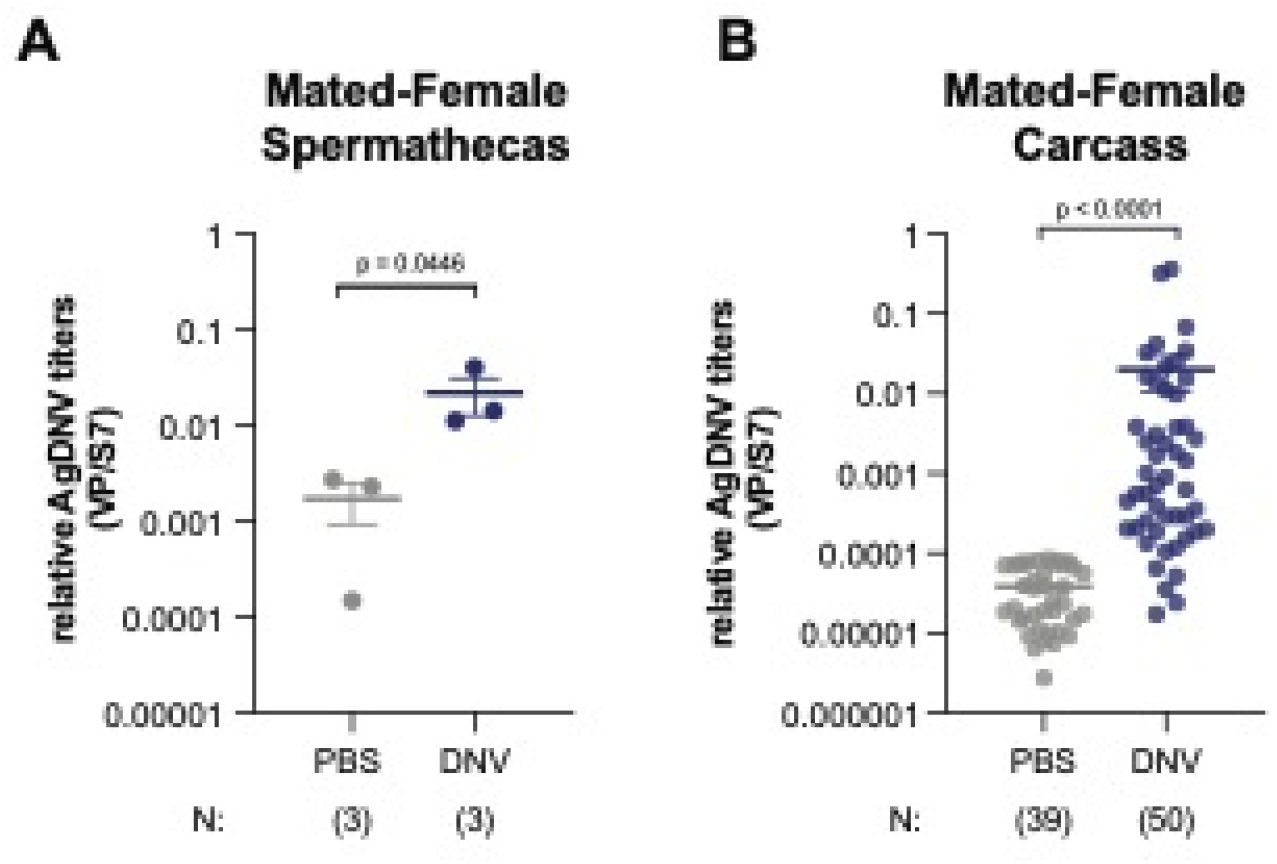
Venereally transmitted AgDNV causes disseminated infection in mated females. (**A-B**) Females mated to AgDNV-injected males have (**A**) detectable AgDNV in the spermatheca (Ln(y) transform, Unpaired Student’s t-test, each point = pool of 5-13 tissues), and (**B**) disseminated AgDNV infection in the carcass (Ln(y) transform, Welch’s t-test, each point = single carcass with spermatheca removed), when analyzed 8-14 days post-mating and relative to females mated to PBS-treated mates. Viral genomes (VP = viral protein) are normalized to host genomes (S7).

### Females venereally infected with AgDNV fail to vertically transmit to F1 progeny

We next assessed whether females sexually infected with AgDNV can vertically transmit the virus to their offspring. Specifically, after females were mated to AgDNV- or PBS-injected males, they were provided a blood meal and an oviposition site. Oviposited eggs were reared to adulthood, and F1 males and females were pooled and assessed for virus. Across all samples tested, AgDNV was not detectable in F1s from either a first or second blood feed (Figure 3A,B). These findings reveal that even though venereally transmitted AgDNV leads to disseminated infection in mated females, they do not vertically transmit to their offspring.

**Figure 3:**
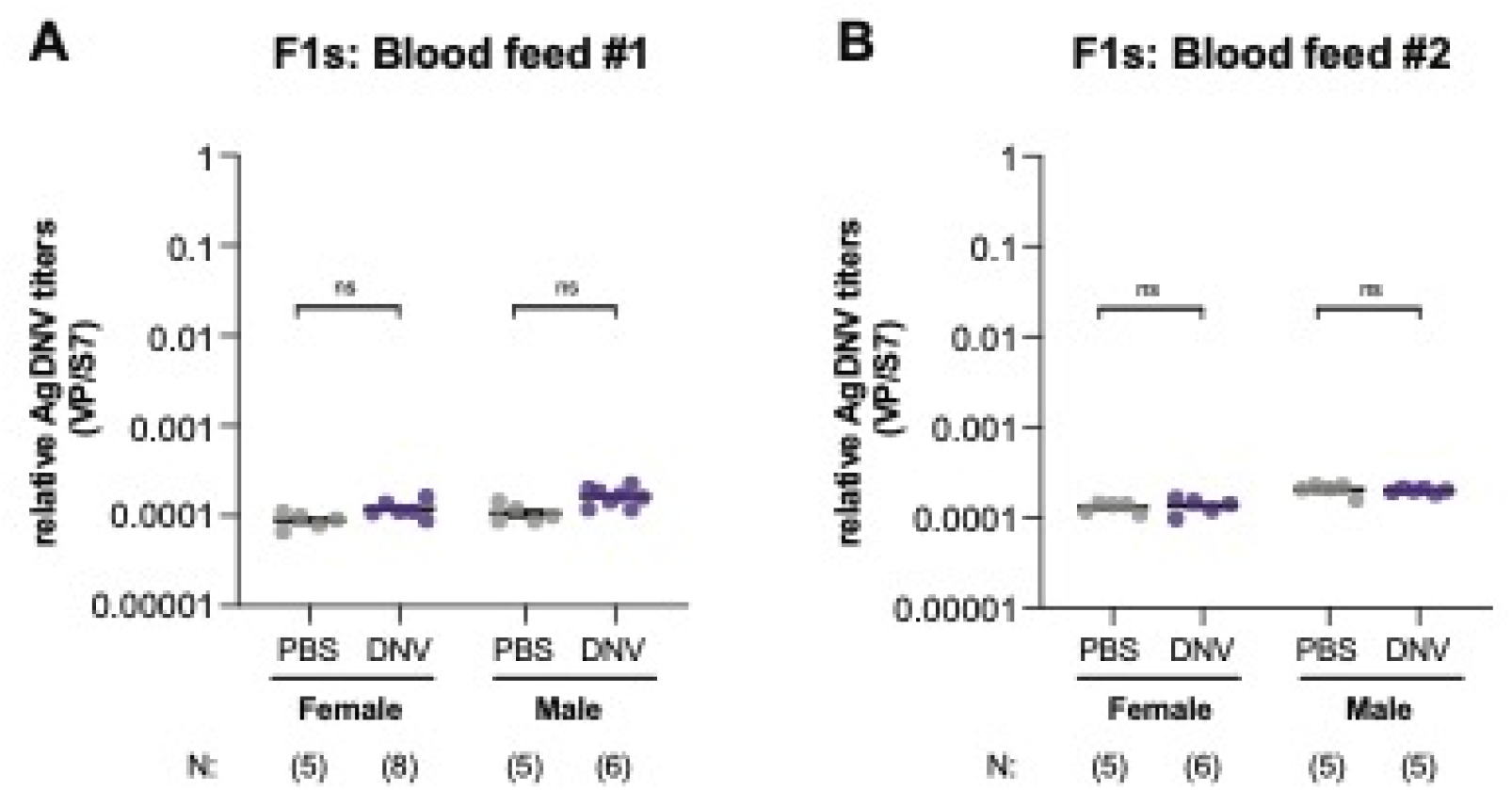
Sexually transmitted AgDNV is not vertically transmitted from mother to offspring. (**A-B**) Females mated to AgDNV-infected males were provided a blood meal and resulting F1s reared to adults. AgDNV was not present in male or female F1s from either a (**A**) first blood feed (One-way ANOVA, Sidak’s correction, each point = pool of 4-5 whole mosquitoes), or a (**B**) second blood feed (One-way ANOVA, Sidak’s correction, each point = pool of 4-5 whole mosquitoes), as compared to F1s with PBS-injected fathers. NS indicates p-value > 0.05. Viral genomes (VP = viral protein) are normalized to host genomes (S7).

## Discussion

Here we report that venereal transmission of AgDNV leads to disseminated infection in *An. gambiae* females. This finding reveals that infected male releases could be an effective way to introduce AgDNV-based paratransgenic tools into field populations of *An. gambiae* females. More specifically, released males harboring a modified AgDNV would mate with wild females and venereally transmit the virus to them. Then, as AgDNV titers build-up in the female post-mating, a viral-encoded effector could interfere with the ability of the mosquito to transmit pathogens like malaria parasites. Male releases have already been shown to be a safe and effective way to deploy biocontrol agents into mosquito populations (14–16), largely because male mosquitoes do not blood feed and thus are not responsible for pathogen transmission. Treatment of larval breeding sites is another means by which ISV-based control tools could be deployed in the field, as was done for the AaeDNV-based larvicide (5, 12). While *An. gambiae* adults can become infected when AgDNV is added to their larval breeding water in the laboratory (1, 7), treatment of larval breeding sites in field settings poses significant challenges. More specifically, *An. gambiae* often lay their eggs in small temporary breeding sites produced by rain fall, making large-scale application of larvae-targeting tools generally impractical for this mosquito species (18). Therefore, infected male releases offers a logistically feasible method to introduce AgDNV-based control tools into wild *Anopheles* populations.

In a previous study, it was shown that AgDNV is transferred to the female spermatheca immediately post-mating (6), but, did not detect significant dissemination in the female carcass. This discrepancy likely arises due to low statistical power in the previous study; elevated DNV was detected in the carcass but it was not statistically significant (6). In addition, females in (6) were assessed immediately after mating, whereas here we assayed for AgDNV in females 8-14 days post-mating, allowing time for the virus to replicate. Given that AgDNV titers are known to rise gradually in adult females (6, 7), it is not surprising that it took a few days for AgDNV titers to increase to significant levels in mated females. In fact, the gradual increase in AgDNV titers over time is one reason this ISV has been proposed as a candidate for a “late-life-acting” insecticide (7, 19).

For infected male releases to be successful, AgDNV must not only be venereally transmitted, AgDNV-infected males must also be fit and able to mate at rates competitive with wild males. Here, we found that AgDNV infection in males does not affect mating rates, suggesting that infected males would be competitive in the field, although mate-choice experiments are still needed to test this directly. We also assessed the tissue tropism of AgDNV in male mosquitoes and found that both the testes and the MAGs become infected, indicating that venereally transmitted virus could be transferred via sperm (produced in the testes), the mating-plug (produced from the MAGs), or both. After mating, sperm is stored in the female spermatheca and the mating-plug is digested in the female atrium (uterus) (20); since we see AgDNV in both the spermatheca and the carcass of mated females, it is likely that both components contribute to sexual transmission of AgDNV.

Although venereally transmitted AgDNV produces disseminated infection in females, we found no evidence of vertical transmission from these females. Interestingly, previous studies demonstrated vertical transmission of AgDNV when both male and female mosquitoes were directly infected by exposure to virus during larval development (1). There are several possible reasons why we did not observe vertical transmission in our experiments here. First, although AgDNV was present in the female body, it may not have infected the ovaries/oocytes. Secondly, AgDNV present in the spermatheca may not be directly associated with sperm cells and therefore not likely to enter the egg during fertilization. Finally, it is possible that AgDNV titers simply were not high enough for vertical transmission to occur in these mosquitoes. Without vertical transmission, an AgDNV-based vector control tool would not spread trans-generationally following infected male releases. This means that a greater frequency of male releases would be required to have an impact on wild *An. gambiae* populations, but on the other hand, this feature may be advantageous from a public acceptance stand-point, because the virus and its encoded effectors would not be heritable across multiple generations. This feature would give more user control over the application of these paratransgenic tools.

Our studies here focused exclusively on the infection and transmission dynamics of wild-type AgDNV. This contrasts with several previous studies of AgDNV-based paratransgenesis that have largely focused on a two-virus helper/transducer system (consisting of one wild-type AgDNV and one defective, GFP-expressing AgDNV) (1, 6, 13). While this helper/transducer approach offers a larger capacity for transgenic cargo, it is far more practical to utilize a single, self-replicating transgenic virus for application in field settings. Viral packaging efficiency limits the size of transgenes that can be encoded in a single AgDNV vector, but small anti-pathogenic effectors or short-hairpin RNAs targeting host processes could be expressed under an endogenous promoter to limit the mosquito’s ability to transmit pathogens. Further engineering and testing are still needed to identify and validate the best effectors for this system.

Finally, besides offering paratransgenic tools, ISVs can also have natural pathogen-blocking effects (21). For example, AgDNV has an inhibitory effect on the emerging arbovirus Mayaro virus (22), which can be transmitted by *Anopheles* (23). Wild-type AgDNV may also inhibit additional pathogens such as the malaria parasite, though this remains to be explored. Together, our findings underscore the importance of basic research into the dynamics of wild-type ISVs such as AgDNV, which hold promise for both basic and applied purposes.

## Conclusions

These findings reveal that if AgDNV is developed into a paratransgenic tool for *An. gambiae* mosquitoes, it could be potentially deployed in the field via infected male releases. *Anopheles gambiae* spreads more fatal infections to humans than any other vector—yet to date—we still lack genetic tools for targeting this species and AgDNV may fill this gap.

## Methods

### Production of AgDNV (GenBank: EU233812.1)

Moss55 cells, which naturally lack wild-type AgDNV (1), were transfected with a plasmid containing the complete wild-type genome of AgDNV, as previously described (1, 6, 13). Briefly, 6-well plates were seeded with ~3 x 10^6 cells per well. The following day, transfections were performed using Lipofectamine LTX with Plus Reagent (ThermoFisher) and 2.5ug of AgDNV-containing plasmid (pAgDNV) per well. Three days after transfection, AgDNV virus was harvested as follows: cells were washed and re-suspended in PBS, lysed via three freeze/thaw cycles (alternating −80°C and room temperature), and then spun down to remove cell debris. Supernatant was collected, aliquoted, and stored at −80°C. This supernatant served as the AgDNV viral inoculum for all experiments.

### Mosquito rearing

Mosquitoes were reared at 27°C with ~80% humidity under a 12h:12h light: dark cycle. Larvae were fed ground Tetramin flakes daily. At the pupal stage, males and females were separated to maintain virgin adults. Adults were provided 10% sucrose solution throughout.

### Microinjections

2-3 days post-emergence, virgin males were inter-thoracically injected with ~200nl of either AgDNV viral stock or PBS (negative control) using a Nanoject II (Drummond Scientific Company) and glass capillary needles back filled with mineral oil. Mosquitoes injected with AgDNV each received an estimated 10^6 vge (viral genome equivalents). Mosquitoes were cold-anesthetized using ice and a cold-block during injections, and immediately placed to recover at room temperature after being injected.

### Mating

When AgDNV- and PBS-injected male mosquitoes were 14 days post-injection (dpi), virgin females were introduced into their cages using a mouth aspirator (John W. Hock). Females were added in a 1:2 ratio with males, and these mixed cages were left for three nights to allow for mating. Afterwards, the mixed sex cage was briefly chilled at 4°C to anesthetize the mosquitoes. Males and females were then separated over ice using a paintbrush and forceps and transferred to new cages.

### Blood feeding and oviposition for F1s

Female mosquitoes were provided with a blood meal 1 and 7 days post-mating. Blood feeding was done using a 37°C water bath, glass membrane feeders, and human blood sourced from BioIVT. Two days after each blood feeding, females were provided with an oviposition site (petri dish with wet cotton and filter paper). Eggs were collected three days later and immediately hatched and reared as described above.

### Tissue collections

Dissected tissues were either stored dry (carcasses) or collected in 50ul-100ul of PBS (spermathecas, testes, MAGs) and stored at −80°C until DNA extraction.

Males: Male reproductive tissues (testes and MAGs) were collected in pools of 12, while male carcasses (body without reproductive tract) were analyzed individually. Males were analyzed at 11, 13, 15, and 18 dpi.

Females: Female mating status was determined by visual examination of the spermatheca for the presence or absence of sperm. Only mated females were further processed for AgDNV quantification. Spermathecas were pooled into groups of 5-13 tissues, while female carcasses (body without spermatheca) were analyzed individually. Females were analyzed at 8-14 days post-mating (estimated, because exact time of mating was not known).

F1s: F1s were collected as whole mosquitoes, and both males and females were analyzed in groups of 4-5 individuals. F1 adults were maintained in a mixed sex cage until collection.

### DNA extraction from mosquito tissues

Tissues were first homogenized using either sterile pellet pestles (Fisherbrand) and an electric pestle grinder (Kimble) (spermathecas, testes, MAGs), or stainless steel beads (OPS Diagnostics) and a TissueLyser II (Qiagen) (carcasses). After homogenization, TL buffer (Omega, Bio-Tek) was added to reach a final volume of 200ul. DNA was then extracted from the samples using the E.Z.N.A Tissue DNA kit (Omega, Bio-Tek) according to manufacturer’s instructions. Proteinase K incubation was carried out for 60min at 55°C in a water bath with vortexing every 15-20min, and final samples were eluted twice with 50ul elution buffer warmed to 70°C. Sample quantity and quality were checked using a Nanodrop (Thermo Fisher) before proceeding to qPCR.

### Quantification of AgDNV

Quantification of AgDNV viral stock produced from transfected cells: One aliquot of virus stock was removed from storage at −80°C and thawed on ice. Using Turbo DNase (Invitrogen) the sample was treated to remove any remaining plasmid DNA, which may be left over from the transfection and would interfere with viral quantification. Next, viral DNA was extracted from the stock using the E.Z.N.A Tissue DNA kit (Omega, Bio-Tek) according to manufacturer’s instructions for cultured cells, starting with Proteinase K treatment. qPCR was then run on a Roto-Gene Q (Qiagen) using PerfeCTa SYBR Green master mix (Quanta-bio) and primers targeting the viral protein (VP) gene on the AgDNV genome. Samples were run alongside a standard curve, consisting of a pAgDNV dilution series, to allow for quantification of viral genome equivalents per ml (vge/ml). VP primer sequences were the same as those used in (22): 5’-GGC ATC AAT GTG GGA CCA AG-3’ (forward) and 5’-CCG TTA GCA AGC GTT GTC TG-3’ (reverse). qPCR cycling conditions were: 95°C for 5min followed by 40 cycles of 95°C for 10sec followed by 60°C for 30sec (data acquisition step). A melt curve analysis was run at the end of cycling by going from 72°C to 95°C with 5sec per step.

Quantifying AgDNV from mosquito tissues: Following DNA extraction, samples were analyzed by qPCR, as described above, except that a pAgDNV standard curve was not run. Instead, primers targeting the host ribosomal protein gene S7 were run alongside primers targeting the viral VP gene. Relative AgDNV titers were then determined by normalizing VP quantification (viral genomes) to S7 quantification (host genomes). This normalizes for differences in the amount of total DNA obtained from different tissue types (e.g., testes vs carcass). S7 primer sequences were: 5’-AAG GGT TGC GTG CTA GTG AA-3’ (forward) and 5’-TAA CGG CTT TTC TGC GTC CA-3’ (reverse). AgDNV dissemination rates in mated female carcasses (text associated with Figure 2B) were estimated by using the highest detectable VP/S7 value found in control female carcasses (0.0000845 VP/S7) as the cut-off for determining AgDNV positive (n = 45, 90%) or negative samples (n = 5, 10%).

### Statistical Analyses

Data for relative AgDNV titers were first tested for normality (Kolmogorov-Smirnov, D’Agostino & Pearson) and then, where appropriate, Ln(y) transformed to achieve normality. When analyzing two groups, unpaired student’s t-tests were used for data with equal variances, and Welch’s t-tests were used for data with unequal variances. If analyzing more than two groups, one-way ANOVA (equal variances) or Brown-Forsythe/Welch ANOVA (unequal variances) was used with either Sidak’s or Dunnett’s multiple testing correction, respectively. Mating and survival rates were analyzed using Fisher’s Exact tests. Statistical tests are also indicated in figure and table legends. A p-value cut-off of < 0.05 was used to determine significance. Analyses were performed with either GraphPad Prism or JMP software.

## List of Abbreviations

AgDNV: *Anopheles gambiae* densovirus
ISV: insect specific virus
MAGs: male accessory glands
pAgDNV: plasmid containing complete AgDNV genome
VP: viral protein (gene)
S7: host ribosomal protein gene
dpi: days post-injection
vge: viral genome equivalents

## Declarations

### Ethics approval and consent to participate

NA

### Consent for publication

NA

### Availability of data and materials

All data generated or analyzed during this study are included in this published article.

### Authors’ contributions

KLW and JLR conceived and designed the study and analyzed the data. KLW, JLR, and HCM wrote and edited the manuscript. KLW performed the experiments. JLR provided materials and reagents. RMJ contributed protocols, reagents, and experimental insights. KLW and HCM designed mosquito graphics. All authors read and approved the final manuscript.

## Acknowledgements

The authors wish to thank Ms Amelia Romo and Ms Kaylee Montanari for their assistance rearing the *An. gambiae* colony mosquitoes.

